# Inference of polyploid origin and inheritance mode from population genomic data

**DOI:** 10.1101/2021.07.19.452883

**Authors:** Alison Dawn Scott, Jozefien D. Van de Velde, Polina Yu. Novikova

## Abstract

Whole-genome duplications yield varied chromosomal pairing patterns, ranging from strictly bivalent to multivalent, resulting in disomic and polysomic inheritance modes. In the bivalent case, homeologous chromosomes form pairs, where in a multivalent pattern all copies are homologous and are therefore free to pair and recombine. As sufficient sequencing data is more readily available than high-quality cytological assessments of meiotic behavior or population genetic assessment of allelic segregation, especially for non-model organisms, here we describe two bioinformatics approaches to infer origins and inheritance modes of polyploids using short-read sequencing data. The first approach is based on distributions of allelic read depth at the heterozygous sites within an individual, as the expectations of such distributions are different for disomic and polysomic inheritance modes. The second approach is more laborious and based on a phylogenetic assessment of partially phased haplotypes of a polyploid in comparison to the closest diploid relatives. We discuss the sources of deviations from expected inheritance patterns, advantages and pitfalls of both methods, effects of mating types on the performance of the methods, and possible future developments.

## Introduction

Polyploid organisms have more than two sets of chromosomes as a result of whole-genome duplication (WGD) and are typically described as belonging to one of two categories: ***auto***polyploids being formed by WGD within a single lineage, and ***allo***polyploids in the case of hybridization between divergent lineages. Though observations of chromosome behaviour during meiosis were initially the only information available for assigning polyploid “types” ***(1)***, we now have to distinguish between this original **cytogenetic**(genetic according to Li et al. ***(2)*** ) classification of auto- or allo-polyploidy, and the now common **phylogenetic** classification (taxonomic according to Li et al., we prefer the term phylogenetic as evolutionary relationships are not always reflected by taxonomy). Furthermore, there is a distinction between inheritance mode (allelic segregation) in polyploids and their classification by either cytogenetic or phylogenetic means.

### Polyploid “types” and inference of inheritance patterns

#### Cytogenetic inferences

Depending on the amount of divergence and structural differences between the duplicated chromosomes, they may or may not pair and recombine during meiosis. Therefore, the origin of polyploidy is traditionally associated with the manner of chromosome segregation. In allopolyploids with sufficient divergence between the subgenomes, only homologous chromosomes (i.e., from the same subgenome; **homeo**logous characterizes the relationship between chromosomes brought together via hybridization among species) would pair and recombine, resulting in disomic inheritance mode ***(3, 4)*** and practically diploid-like behavior. In autopolyploids, where each chromosomal version is homologous to each other, chromosomes show a random assortment during meiosis and can form bivalents or multivalents ***(5, 6)***. In, e.g., a tetraploid with four identical subgenomes (both a cyto- and phylo-genetic autopolyploid), pairing can occur among any of the four homologous chromosomes during meiosis, either via formation of multivalents or random bivalents. Therefore, gametes formed by this polyploid can contain any combination of its four alleles, as preferential pairing is absent. In a cytogenetic allopolyploid, bivalents are formed among homologous chromosomes, while pairing of homeologous chromosomes is prevented (e.g. pairing only happens *within* subgenomes, not between or among). Thus the types of gametes formed by an allopolyploid are disomic, and the allelic combinations within them are limited compared to those formed by an autopolyploid.

As is often the case in biology, real-life examples deviate from the binary extremes and can be realized anywhere along a spectrum from bivalent pairing with disomic inheritance to fully multivalent pairing and polysomic modes. For example, autopolyploids initially form multivalents following their origin, but eventual diploidization will result in strictly bivalent pairings. Similarly, allopolyploids may exhibit primarily bivalent pairing due to divergence among parental genomes, but retain homology of some chromosomes thus yielding multivalents.

#### Phylogenetic inferences

With the increased availability of sequence data, the origins of countless polyploid organisms have been classified (e.g., auto, allo, or some combination thereof) via phylogenetic inference ***(7)***. While cytogenetic classification is based on chromosome pairing behavior (e.g., formation of either bivalents or multivalents), phylogenetic classification of polyploids is based on the relatedness of a polyploid’s constituent subgenomes in a tree. That is, if homeologous gene copies within the polyploid taxon are more closely related to each other than to gene copies from another taxon, it is commonly classified as an autopolyploid. Conversely, the homeologous gene copies of a phylogenetic allopolyploid would not form a clade, rather one or more homeologs would be more closely related to another taxon than to other homeologs.

Generally, phylogenetic inferences of polyploids are focused on identifying their origin, either from a past hybridization event or a duplication within a lineage. Unlike cytogenetic state and inheritance mode, the phylogenetic context in which a polyploid emerged cannot change. However, it may become harder to infer phylogenetic origin with the passage of time and accompanying genome evolution, lineage extinction, etc.

#### Segregation patterns in offspring & gametes

To study inheritance patterns, the gold standard is to observe genotypes in parents and their offspring. Observed inheritance patterns in parents and their F_1_ hybrids can then be compared to the different possible inheritance modes, e.g., disomic or tetrasomic inheritance ***(8)***. However, such an approach is not always feasible for a number of reasons, including the time and expense involved in breeding studies. One possible “shortcut” is single-cell sequencing of gametes ***(9)***, which permits high throughput genotyping on many thousand haplotypes and can easily be expanded to polyploid taxa ***(10)***. While phylogenetic inference of polyploids is typically used to understand something that happened in the past, namely the origin of a polyploid, understanding the current process of inheritance allows insights into the ongoing evolution of polyploid taxa.

#### Conflict between inferences

When making inferences about the origin of a given polyploid, it is possible to observe, e.g., a cytogenetic allopolyploid that is a phylogenetic autopolyploid, and other seemingly disparate combinations. For example, autotetraploid *Arabidopsis arenosa* has a tetrasomic inheritance mode despite forming bivalents during meiosis ***(11)***. The possibility of different observations from a) phylogenetic classification of polyploid origin, b) cytogenetic classification of polyploid meiosis and c) polyploid inheritance patterns highlights the importance of integrating multiple inference methods to understand polyploid evolution. The origin of polyploids often dictates the way we tend to analyse genome sequencing data. Sequencing data from allopolyploids with sufficient divergence between subgenomes can often be split either by mapping to the combined reference of the parental genomes with subsequent filtering for primary alignment in proper pairs ***(12, 13)*** or by using SNP markers specific for each parent ***(14, 15)***. Divided subgenomes are then analysed as if they were separate taxa. Autopolyploid data are usually mapped to a single reference genome and the number of alleles called at each site is equal to ploidy ***(16, 17)***, similar to pooled sequencing experiments, where number of alleles corresponds to number of haploid individuals (see also Bohutinska et al. chapter). The inheritance mode of polyploids impacts the evolutionary inferences we draw from analysing such data and deviations from the assumed mode can strongly bias the estimates of genetic diversity and population divergence ***(18)***.

### Sources of mixed inheritance patterns

#### Homeologous exchanges

Auto- and allopolyploidy origins do not have a strict correspondence in chromosomal pairing behavior. Although rarely, the homeologous chromosomes from the different subgenomes in allopolyploids can sometimes pair and recombine ***(19–21)***. Homeologous exchanges are common in recent allopolyploids, for example in *Tragopogon miscellus **(22)***, cultivated peanut ***(23)***, and resynthesized allotetraploid rice ***(24)***. In more established allopolyploids, homeologous exchanges are rare and appear to be under genetic control. For example, in allohexaploid wheat, *Ph1* locus ***(25–27)*** carries the major crossover gene *ZIP4* and prevents homeologous crossovers. *PrBn* locus in oilseed rape (*Brassica napus*) has a similar effect ***(28)***. Preferential homeologous pairing in *A. suecica* also has a genetic basis ***(29)*** (*BOY NAMED SUE* locus) but also appears to be regulated on the transcriptional level of meiotic genes ***(12)***. This suggests that homeologous exchanges are mostly deleterious and allopolyploids do not only rely on the divergence levels between the subgenomes to prevent these generally undesirable genome reshufflings. Despite stricter regulation of preferential pairing between homologs, homeologous exchanges do happen, and we expect such regions to show apparent mixed inheritance patterns.

#### Rediploidization

Rediploidization is the process by which a polyploid gradually behaves more and more like a diploid - e.g., a) multivalent formation is replaced by strictly bivalents and b) polysomic inheritance is no longer observed ***(4, 30, 31)***. In general, the term rediploidization is used when discussing the evolution of autopolyploids, as allopolyploids often behave largely as diploids (cytogenetically speaking). Unlike a whole-genome duplication event, which happens quickly and is followed by a polyploid establishment process, rediploidization takes place over longer periods of time - anywhere from 20-30 generations ***(32)*** to tens of millions of years ***(33)***. Furthermore, as rediploidization is an ongoing process, it is entirely likely that different lineages are experiencing the process in different ways ***(33)***, such that a handful of observations may not represent the true evolutionary dynamics at play. For example, when bivalent and multivalent pairing are both possible, different chromosomes and lines may exhibit an inconsistent pattern ***(34)***. It is possible that parts of the genome show disomic inheritance, whilst other regions do not, the latter associated with, e.g., interchromosomal translocations and inversions. This drawn out rediploidization process can therefore create mixed inheritance patterns when we observe polyploid taxa caught “in limbo” between an autopolyploid and a fully diploidized state.

#### Interspecific introgression

Interspecific introgression can become another source of mixed inheritance patterns. Recent studies in *Arabidopsis* plants ***(35, 36)*** and *Neobatrachus* frogs ***(37)*** suggested that despite the apparent isolation of the diploid species, tetraploid lineages in those groups can hybridize. This suggests that polyploidy can mediate interspecific gene flow ***(38)***. One explanation of this phenomenon in seed plants is based on the hypothesis that polyploidy permits breakdown of endosperm-based incompatibility, thus lifting the postzygotic barrier and allowing hybridization among species ***(39, 40)***. Another explanation could be the more effective masking of recessive hybrid incompatibility loci in polyploids. Hybridization can happen (1) between tetraploids with different ancestors and (2) between a tetraploid and an unreduced gamete of a non-ancestral diploid species. If genomes of both species are sufficiently divergent to prevent homologous pairing, both cases would result in mixed inheritance patterns.

Except for lab-generated polyploids, we are not privy to the polyploidization events of the taxa we study. Furthermore, researchers rarely have access to cytogenetic, genomic, and breeding resources to allow direct observation of chromosome pairing, phylogenetic context, and inheritance patterns of their study systems. The methods we describe here are meant to facilitate the study of polyploids and the consequences of their whole genome duplications, using bioinformatic approaches to infer origin and inheritance modes from population genomic data. The first method uses allelic depth distributions within an individual, and classifies their fit to expected distributions for disomic and tetrasomic inheritance modes. The second method leverages haplotype distances along the genome to infer origin of polyploidy, and deviations from expected inheritance modes.

## Materials

The computational inference of polyploid inheritance mode requires sequenced datasets of the polyploid and related diploid lineages. We demonstrate both approaches using a reference genome and whole-genome resequencing data with short paired reads, however, these methods can also work with reduced representation (e.g., RADseq) or exome-capture sequencing data ***(37)***. Data from several individuals per lineage (we use at least 4 individuals in our examples) will account for within-lineage variation (but not deep coalescence) and ensure the robustness of the approach. Analysis of the whole-genome sequencing data usually requires a high-performance computer, however, the computational load massively depends on the genome size and sequencing depth. To illustrate the approaches we describe in this chapter, we used previously published ***(13, 41–43)*** raw reads from *A. suecica* (SRR2084158, SRR3123767, SRR3123760, SRR2084157), *A. arenosa* (SRR2040811, SRR2040813, SRR2040820, SRR2040824, SRR3111444, SRR3111445, SRR2040800, SRR2040802) and *A. thaliana* (SRR519491, SRR519660, SRR519519, SRR519685, SRR519656, SRR519575, SRR519494, SRR519484, SRR519647, SRR519714, SRR519613, SRR519641, SRR519684, SRR519505, SRR519489, SRR519687).

### Methods

**The first approach** to infer inheritance modes of sequenced samples is based on the differences in allelic read depth expectations. In a diploid sample, the distribution of allelic depth at the biallelic heterozygous (Aa) sites is expected to be centered around 0.5, where 50% of the reads belong to the first (A) and another 50% - to the second (a) allele. In tetraploid samples, biallelic sites can form three heterozygous genotypic states: AAAa, AAaa, Aaaa. The expectations of the relative frequency of each genotype in tetraploids are different for autotetraploids (with tetrasomic inheritance) and allotetraploids (with disomic inheritance) ***(11, 44)***; (see Note 1). In the case of an allotetraploid origin or a long evolution under a disomic inheritance of an original autotetraploid, we expect to see an excess of AAaa genotypes in such an individual. Even with ongoing rediploidization (and accompanying copy number reduction), the distribution would still permit inference of inheritance mode. Similarly, on a population level, we expect to see an excess of SNPs with intermediate allele frequencies. The amount of such excess depends on the evolutionary history of the polyploid and its diploid relatives and can be compared to simulated data or computationally combined diploid data into tetraploids of different origins. In order to illustrate the method, we show examples of allelic depth distributions from both simulated and real sequencing data (Fig. 1).

**Figure 1.**
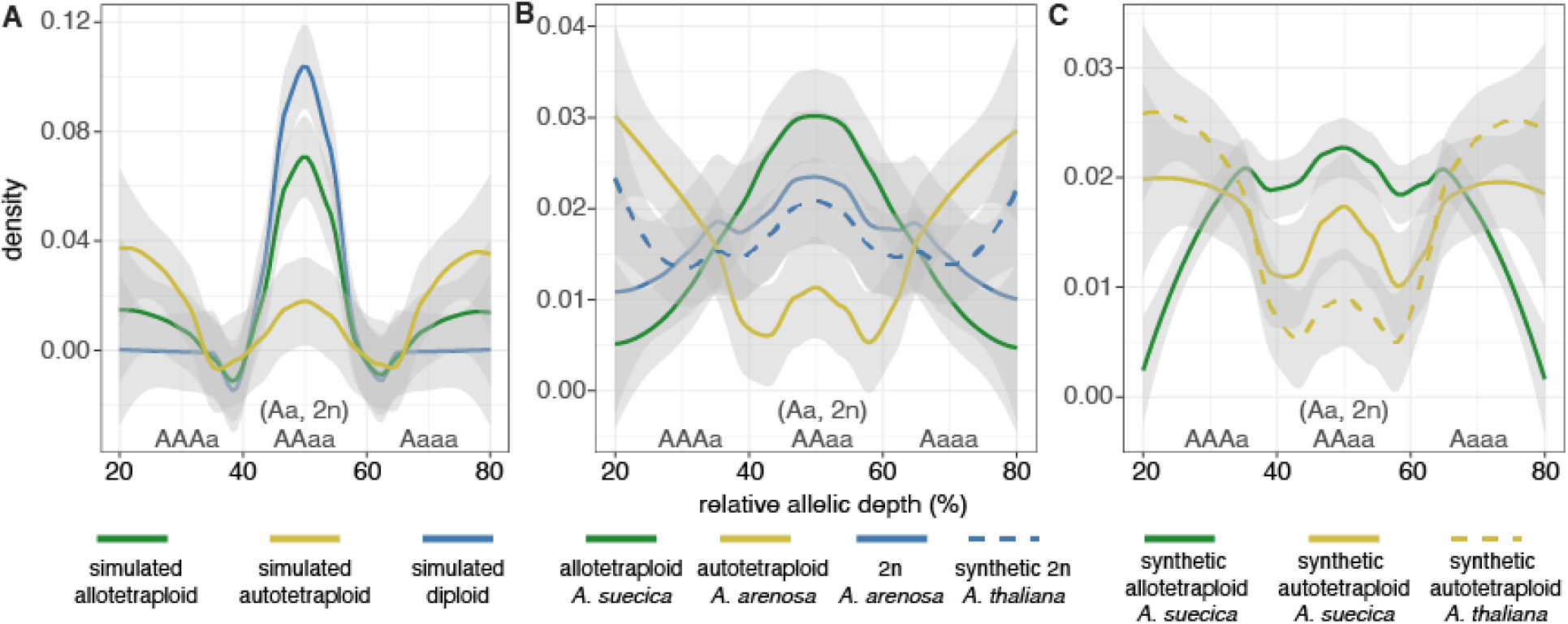
Inference of inheritance modes of polyploids using allelic read depth distributions. **a**) Simulated diploid (blue line) and allotetraploid (green line) samples show excess of intermediate ratios of allelic depth: Aa or AAaa genotypes, respectively. This contrasts with simulated autotetraploid (yellow line), where the intermediate allelic ratio does not have such a pronounced peak. **b**) Real data examples showing the distributions of allelic read depth for diploid *A. arenosa* (blue line), autotetraploid *A. arenosa* (yellow line) and allotetraploid *A. suecica*. Because *A. thaliana* is highly inbred due to selfing, we combined sequencing data of two different accessions to show expected distribution of an “outcrossing” diploid *A. thaliana* (dashed blue line). **c**) Synthetic controls for the real data examples. Green line - computationally combined sequencing data of two haploid *A. thaliana* accessions and one diploid *A. arenosa* make an ‘outcrossing’ allotetraploid with a similar genetic composition to *A. suecica*. Yellow lines show synthetic autotetraploids: dashed - four combined accessions of *A. thaliana*, solid - two combined accessions of diploid *A. arenosa*.

We first applied our method to simulated diploid and polyploid individuals (Fig. 1a). Briefly, we simulated a tree, genome, and sequences with Zombi ***(45)***. These sequences were then used to simulate Illumina HiSeq reads using InSilicoSeq ***(46)***. Based on tree topology, we selected eight diploid individuals and one reference accession. The diploid genomes were combined to form four autotetraploids and four allotetraploids, mimicking the immediate outcome following polyploidization. These simulated accessions were then analyzed as described for the real data below. The obtained distributions of allelic ratios for the simulated data follow the expected pattern: diploid and allotetraploid samples have an excess of intermediate ratios (Aa and AAaa), when genotypes of the simulated autotetraploids are more evenly distributed.

As a real data example, we chose *Arabidopsis* species with well-described evolutionary histories and known inheritance modes ***(11–13, 41, 42, 47)***: *A. thaliana* - a diploid, *A. arenosa* - both diploid and autotetraploid lineages, and *A. suecica* - an allotetraploid hybrid between *A. thaliana* and *A. arenosa*. This example also demonstrates the importance of taking a mating type of the species into account when estimating inheritance modes (see Note 2), as this method is based on the heterozygous sites only - selfing species will produce an excess of homozygotes compared to outcrossers. Both *A. thaliana* and *A. suecica* are self-compatible and mostly selfing plants, when both diploid and tetraploid lineages of *A. arenosa* are obligate outcrossers. For our example, we mapped 12 different *A. thaliana*, 4 diploid and 4 tetraploid *A. arenosa* lineages, and 4 *A. suecica* accessions to the *Arabidopsis lyrata* reference genome ***(48)***. For mapping, we chose the bwa mem algorithm ***(49)***, which allows split reads and is therefore suitable for cross-species reference mapping with bigger divergence times. For this comparative analysis between different species, we chose only the genic regions which are more conserved and usually better covered compared to the intergenic regions. Apart from increasing the average quality of the SNPs, such an approach of region filtering also substantially decreases the computational time for further analysis. Distribution of allelic depth at biallelic SNPs per individual can be obtained in different ways (for example, by parsing the output of samtools mpileup ***(50)*** or GATK ***(51)***); we chose nQuire software ***(52)***, which was originally written for estimation of ploidy levels. Using the “create” command of nQuire, we first generate a binary .bin file from the bam file. Then we apply the “denoise” algorithm implemented in nQuire to scale down the noise present from low-quality SNPs and potential misalignments. And finally, we visualise the base frequencies for each individual using the command “histo” from nQuire, which creates a histogram in ASCII format, easily parsed in R ***(53)***.

The real data example of the distribution of allelic depth follows the expectations: the obligatory outcrossing *A. arenosa* diploid lineage (Fig. 1b, blue line) has the highest peak at the 50% allelic ratio (Aa), while in the autotetraploid *A. arenosa* (Fig. 1b, yellow line) AAAa and Aaaa genotypes seem to be more common. In the case of *A. thaliana* species, we need to consider that it is mostly a selfing plant, resulting in a dearth of heterozygous sites and therefore no peak (at genotype Aa). To show the expected patterns based on *A. thaliana* sequences, we combine reads from two different *A. thaliana* accessions to obtain a “synthetic” diploid *A. thaliana* (Fig. 1b, dashed blue line), which follow a similar pattern to a diploid outcrossing *A. arenosa*, except for the tails of the distribution. This difference in the tails, where the synthetic diploid *A. thaliana* more closely resembles true tetraploid *A. arenosa*, may be explained by rare heterozygous sites within *A. thaliana* accessions. *A. suecica* (Fig. 1b, green) is also a selfer and an allotetraploid with an extremely high divergence (about 5My or 9%) ***(41, 54)*** between the subgenomes: the excess of intermediate alleles (AAaa genotype) is prominent.

Although in these examples (Fig. 1b) the differences in the distributions are obvious, in order to make the inferences we still prefer to compare the real data examples to the synthetic combinations of the real data forming synthetic auto- and allo-tetraploids from the exact diploid relatives. We formed synthetic autotetraploid *A. arenosa* by combining reads from two different outcrossing diploid *A. arenosa* individuals (Fig. 1c, solid yellow line) and formed a synthetic autotetraploid *A. thaliana* by combining four selfing *A. thaliana* accessions (Fig. 1c, dashed yellow line). Both of those distributions do not show an excess of intermediate allelic ratio genotypes (AAaa) and resemble real autotetraploid *A. arenosa* data (Fig. 1b, yellow line). We have also combined two “haploid” selfing *A. thaliana* and one diploid outcrossing *A. arenosa* to simulate an outcrossing allotetraploid (Figure 1c, green). This distribution shows an excess of intermediate frequencies, but also differs substantially from the real *A. suecica* data, because *A. suecica* species is selfing and all the biallelic sites are distributed around at 50% allelic ratio and represent the differences between the subgenomes (AAaa, where AA - *A. thaliana* and aa - *A. arenosa* variant, for example). In cases where the differences are not as obvious, one could use, for example, Mann–Whitney *U* test to compare the distributions of rations between intermediate (40–60%) and rare (<30%) allele frequencies in real data samples and synthetically combined controls.

The drawback of the approach based on allelic frequencies is that any deviation from the inheritance mode at the origin of the polyploid (for example, due to ongoing introgression with other species, re-diploidization, or homeologous exchanges) will lead to an averaged mixed signal on the relative allelic depth distributions. Inspecting inheritance mode along the genome, in this case, would provide a more reliable result and pinpoint the regions leading to the mixed signals. Next, we describe an approach that allows investigating inheritance mode along the genome using the same type of input data.

**The second approach** uses distance-based phylogenetic relatedness between haplotypes. Ideally, if one had a full high-quality assembly of all four chromosomal copies of a tetraploid and closely related diploid species, the identification of the origin, evolutionary history, and inheritance mode of such tetraploid would be a matter of inferring phylogenetic relationships between each tetraploid allele and the diploids. While this sounds straightforward, phylogenetic inference over many thousands of loci is complicated by, e.g., poor resolution within regions and considerable conflict among regions. Despite substantial advances in long-read sequencing, it is still most common to obtain a long-read-based reference and several short-reads-based re-sequenced genomes. Short reads do not allow a complete chromosome-scale haplotype-phased resolution, however, it is possible to distinguish between the two most divergent haplotypes within relatively short and well-covered loci using paired-end information. For example, the phasing algorithm implemented in samtools phase ***(50)*** splits a .bam file resulting from the mapping of short reads to a reference genome into the two most divergent haplotypes (which makes the method suitable only for tetraploids, but not higher ploidy level samples, Note 4). Here, one should be aware that regions with low coverage or a low SNP density would interrupt continuous phasing and randomise haplotypes in the two resulted files, so the phasing of the haplotypes is typically only reliable within relatively short loci. Nevertheless, this approach allows estimation of phylogeny within, e.g., genic regions and has been applied to infer the origins of, for example, allotetraploid *Medicago* species ***(7)***.

Using the same *Arabidopsis* samples as in the first approach (*A. thaliana* - 2n = 2x, selfing; *A. arenosa* - 2n = 2x and auto 4n = 4x, outcrossing; and *A. suecica* - allo 4n = 4x, selfing), we demonstrate the framework to infer inheritance modes using partially (see Note 5) phased haplotypes. Again, we use the synthetic controls for auto and allotetraploids (or tetrasomic and disomic inheritance modes in this case): computationally combined *A. thaliana* and *A. arenosa* samples within and between the species, respectively (Fig. 2a). We describe the procedure starting from the mapped short reads of all the samples to the same reference genome (in our case - *A. lyrata* reference genome ***(48)***):

1. We split the .bam files of the tetraploids using samtools phase ***(50)*** into two .bam files - this step includes both real data polyploids and synthetic controls (Note 5, Figure 2c).
2. Then, we call variants from all the diploid samples and the two phases of the polyploids separately, using the GATK Haplotype Caller, with setting sample ploidy to 2 ***(51)***. To speed up the process we only call the variants on the genic regions, which we assume would also be better covered and therefore better phased. For an estimate of appropriate coverage for SNP calling, ***(55)*** recommend 15x.
3. Then we combine the variants and perform joint-genotyping using two subsequent steps of CombineGVCF and GenotypeGCVF from GATK. For further analysis we filter for biallelic single-nucleotide polymorphisms (SNPs), which are present in all the diploid samples.
4. We transform the variant table to numeric values, by summing up the amount of non-reference alleles and dividing the result by the number of alleles. So, the diploid samples and split tetraploid samples with reference allele would have 0 (AA), non-reference homozygous allele 1 (aa) and heterozygous allele 0.5 (Aa).
5. Then in R ***(53)***, we identified biallelic SNPs present in all 16 diploid samples and encoded them on a scale from 0 (homozygous reference) to 1 (homozygous alternate) in a separate table for each gene. To determine genetic distance to a real or synthetic polyploid, we added one phase from the polyploid to each gene containing at least 5 SNPs. We then calculated absolute pairwise distance between all pairs of samples in each gene variant table that contained all diploids and one phase of one polyploid.
6. We calculate absolute pairwise distance between all the samples in the table, which includes all the diploid individuals and one phase of a polyploid sample, using the standard “dist” function in R with the method set to “manhattan”. If calculated distance between the polyploid sample and any of the diploids equals 0, which can happen with low diversity species (as *A. thaliana* and *A. suecica*) and result in a polytomy, we check that the clonal samples are indeed coming from the same species (list of tip names contains only one species name) and assign this species to this phase of the gene.
7. In the rest and majority of the cases, we then calculate a neighbor-joining tree using the ‘ape’ package in R ***(56)***, where we choose the node right above the sample of interest (one of the polyploid phases) and list all the tips, which descend from this node (see Note 6). In case all the other tips (diploid samples) belong to one species, we assign that phase to that species, in case they include multiple species (both *A. arenosa* and *A. thaliana*), we can not assign a closest diploid and therefore call it “NA”. We illustrate the obtained results for the example data on Fig. 2b. The final results were plotted with the use of karyoploteR ***(57)***.

**Figure 2.**
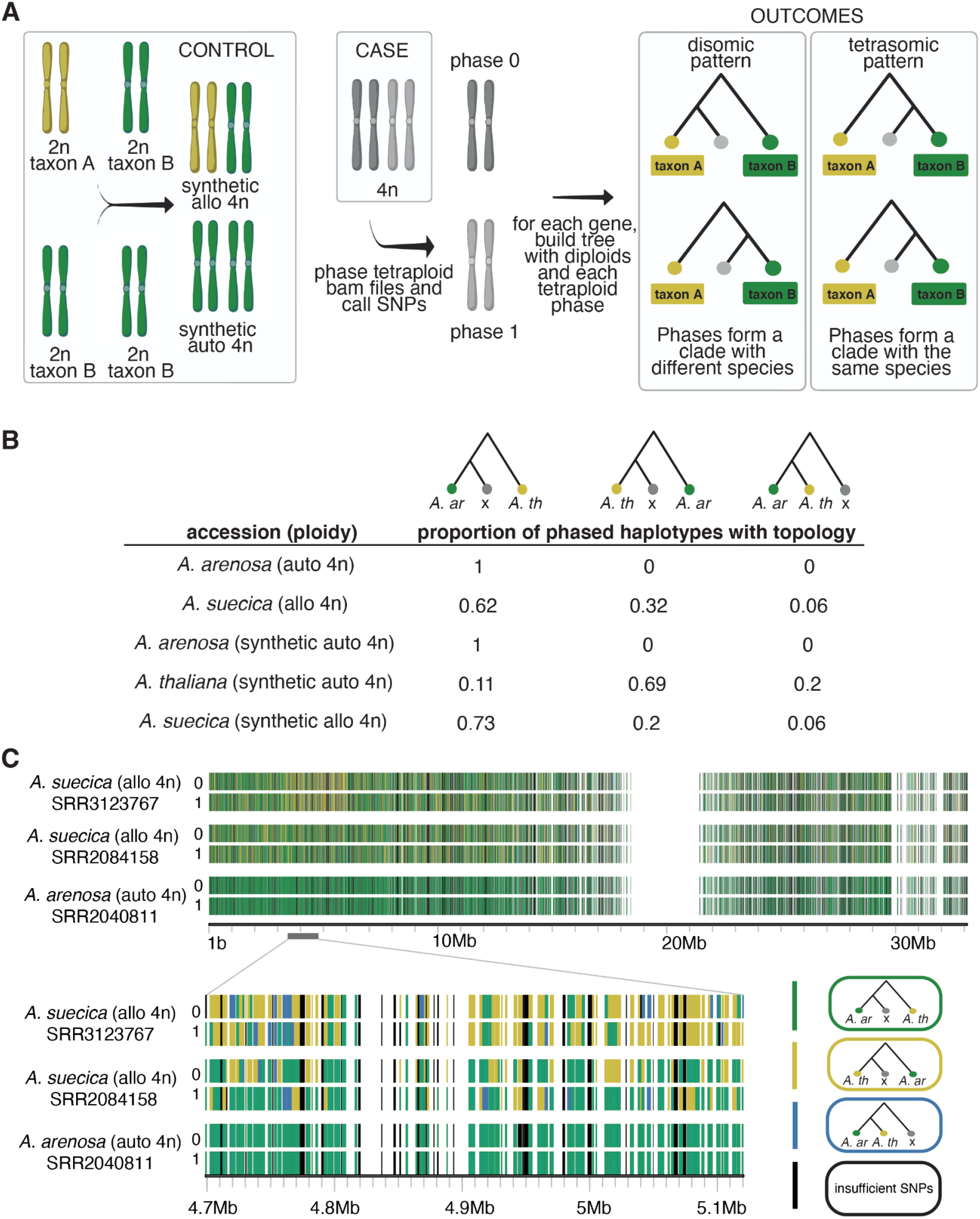
Inference of origin and inheritance modes of polyploids using haplotype phylogenies. **a)** Schematic representation of the computational experiment; created with BioRender.com. Synthetic controls are based on combinations of real sequencing data of the related diploid species (same as in the first approach, Fig. 1c). To obtain a synthetic autotetraploid, we combine two real diploid samples within the same species; to obtain a synthetic allotetraploid, we combine diploid samples of different species. Diploid samples, synthetic controls and the real data from polyploids are mapped to the same reference genome and then real and synthetic polyploids are split into two phases. Assuming that phasing is correct within the genic regions, we investigate the phylogenetic placement of each polyploid phase within each gene/locus: the two phases from the allopolyploid samples would cluster with different diploid species, when the two phases of autopolyploid species would cluster with the same species. **b**) Results obtained from the data of natural *A. suecica* and *A. arenosa* polyploids, as well as synthetic controls. Schematic table represents the average proportions of the genic regions assigned to *A. arenosa*, *A. thaliana* or “NA”, topologies illustrating the assignment procedure are shown above each column. **c**) Assignment of each gene in each phase for two natural allotetraploid *A. suecica* and one natural autotetraploid *A. arenosa* individuals, with scaffold distribution of tree topologies. The zoomed in region shows the homeologous exchange pattern on one of the *A. suecica* accessions (SRR3123767).

From 5,226 genes annotated on the first scaffold of *A. lyrata* reference, on average for each investigated polyploid sample 3763 genes satisfied our filtering criteria. In Figure 2b we provide the average proportion of these genes assigned to one or the other lineage for the natural *A. suecica* and *A. arenosa* polyploids and the synthetic controls. Genes from both synthetic and natural polyploid *A. arenosa* were always assigned to *A. arenosa* lineage, while synthetic *A. thaliana* still had a low proportion of incorrectly assigned genes to *A. arenosa*. This bias towards *A. arenosa* can be explained by about five times closer genetic distance ***(54)*** between the reference *A. lyrata* and *A. arenosa* species compared to the distance between *A. lyrata* and *A. thaliana*. As expected for an allotetraploid, the genes from synthetic and natural *A. suecica* were more evenly assigned to both of the parental (*A. thaliana* and *A. arenosa*) lineages, however, with a certain bias towards *A. arenosa*.

The second method also allows investigation of deviations from the expected pattern of inheritance mode along the genomes of each polyploid sample. The sources of such deviations are mentioned in the introduction (e.g., ongoing rediploidization, interspecific introgression and homeologous exchanges between subgenomes of an allotetraploid). Again, the deep coalescence would resemble inheritance deviation signature, however, it is independent from allelic segregation. This is clearly demonstrated in the case of *A. suecica*, a known allopolyploid (hybridization between *A. arenosa* & *A. thaliana*) which we expect to exhibit preferential (bivalent) chromosome pairing and thus disomic inheritance. Figure 2c shows a stretch of chromosome 1 with a different inferred inheritance mode than the rest of the genome; this is a previously described ***(12)*** homeologous exchange between the first chromosome of *A. suecica* (*A. thaliana* subgenome) and the sixth chromosome of *A. suecica* (*A. arenosa* subgenome) in SRR3123767 (AS530) accession. This region of almost 16Mb of the *A. arenosa* subgenome has been exchanged for the *A. thaliana* copy. Despite this apparent deviation, we would still infer disomic inheritance is the norm for this accession, as this homeologous exchange is not expected to break the previously established preferential pairing among chromosomes. However, this example merits some attention as over time, allopolyploids can accumulate extensive homeologous exchange, resulting in “candy cane” chromosomes (where the closest relative of each stretch of chromosome switches between progenitors). Furthermore, when diploidization follows significant homeologous exchange, the resulting genome will contain a mixture of genomic regions from both parental species. This case highlights the importance of evolutionary history context for the correct interpretation of the observed phylogenetic patterns of the haplotypes along a polyploid genome. In contrast, the first method we describe is more flexible, as it can be applied even when information about polyploid origin is unclear, e.g., when parental genome donors are extinct, unidentified, or otherwise unavailable for sequencing.

### Outlook

The stronger the technology, the less algorithmic effort it requires to interpret, at least in biology. With currently available long read sequencing, assembly of chromosome-scale (or at least substantially longer haplotypes) can replace the need for genomic phasing. This, of course, does not nullify the phylogenetic approach to inheritance mode inference, rather makes it applicable to systems with higher ploidy levels. Long read sequencing may eventually be universally available, but in the meantime short-read based approaches are still useful. Because long-read technology requires high quality, high molecular weight input DNA, this method is inaccessible in some cases, for example, ancient or degraded samples, or those stored in formalin. At the same time, the affordability of sequencing already allows the assesment of hundreds of offspring samples or even gametes to directly measure inheritance patterns, avoiding the estimations altogether. Limitations of this approach again include ancient samples and extinct species, as well as species which are unfeasible to cross (e.g., due to generation time, difficulty of cultivation) and which are hard to obtain in the field, protected species, etc.

### Notes

1. In the method based on allelic depth distributions, we assume that polyploid samples are mapped to a single ancestral (parental) genome, even in the case of allotetraploids, where one of the sub-genomes could be closer to the reference genome in this case. In the case of mapping an allotetraploid to a reference, containing both of the subgenomes, we expect to see a diploid-like allelic depth distribution.
2. When the mating-type is unclear, it is useful to start with estimations of overall heterozygosity levels. Additionally, one should be aware that an outcrossing diploid and a selfing allotetraploid can show a similar distribution of allelic depth at the heterozygous genotypes with a single peak at 50%. In order to distinguish between those types, we recommend including a genome size estimation into the analysis, which can also be done bioinformatically, analysing k-mer frequencies from the same type of input - whole-genome short-read sequencing data ***(58, 59)***.
3. Regarding our second method, it is important to consider the expected variation in topologies among gene trees - different loci will tell different stories, so care must be taken to include many gene trees. Furthermore, the methods described herein do not account for uncertainty in gene tree estimation, nor do they attribute discordance along a chromosome to deep coalescence.
4. The phasing algorithm ***(60)*** implemented in the samtools phase ***(50)*** is tailored to assemble only two “best” haplotypes, which makes it applicable for diploids and tetraploid samples, but probably not higher ploidies. Even with the tetraploid samples, one should be aware that such phasing is only partial. That is why we describe the synthetic controls to use for the correct interpretation of the results.
5. One should be aware that samtools phase will split one bam into two separate files, which do not represent two completely phased chromosomes, but rather short stretches of phased data mixed between the chromosomes (this is also demonstrated in our results, Fig. 2c). The length of the phased stretches will depend on the length of the short reads, the coverage depth, mapping quality in that region and the amount of polymorphisms, which allow to assign the haplotypes into two separate phases. The phasing will be interrupted with the gap between the markers longer than a fragment length.
6. Here we employ distance-based phylogenetics to estimate Neighbor-Joining trees, which is a fast process that scales well to genome-wide analyses. Other phylogenetic inference methods (maximum likelihood, Bayesian inference) are also suitable, but will be more computationally intensive.
7. In the case of method 2, branch lengths are ignored but could be used for an estimate of divergence time between subgenomes. This would require tree estimation by a different method (maximum likelihood, Bayesian inference) and therefore would also require base call information to incorporate a substitution model.

## Acknowledgements

We thank Robin Burns, Filip Kolář and Matt Johnson for their insights and advice during the preparation of this chapter.

